# Targeting RET Kinase in Neuroendocrine Prostate Cancer

**DOI:** 10.1101/622415

**Authors:** Halena VanDeusen, Johnny R. Ramroop, Katherine L. Morel, Anjali V Sheahan, Zoi Sychev, Nathan A. Lau, Larry C. Cheng, Victor Tan, Zhen Li, Ashley Petersen, John K. Lee, Jung Wook Park, Rendong Yang, Ilsa Coleman, Owen Witte, Colm Morrissey, Eva Corey, Peter S. Nelson, Leigh Ellis, Justin M. Drake

## Abstract

Increased treatment of metastatic castration resistant prostate cancer (mCRPC) with second-generation anti-androgen therapies (ADT) has coincided with a greater incidence of lethal, aggressive variant prostate cancer (AVPC) tumors that have lost androgen receptor (AR) signaling. AVPC tumors may also express neuroendocrine markers, termed neuroendocrine prostate cancer (NEPC). Recent evidence suggests kinase signaling may be an important driver of NEPC. To identify targetable kinases in NEPC, we performed global phosphoproteomics comparing AR-negative to AR-positive prostate cancer cell lines and identified multiple altered signaling pathways, including enrichment of RET kinase activity in the AR-negative cell lines. Clinical NEPC and NEPC patient derived xenografts displayed upregulated RET transcript and RET pathway activity. Pharmacologically inhibiting RET kinase in NEPC models dramatically reduced tumor growth and cell viability in mouse and human NEPC models. Our results suggest that targeting RET in NEPC tumors with high RET expression and may be a novel treatment option.

**Statement of Significance:** There are limited treatment options for patients with metastatic aggressive variant prostate cancer and none are curative. Here we identified aberrantly activated RET kinase signaling in multiple models of NEPC. Inhibiting RET restricted tumor growth, providing a novel approach for treating NEPC.

## Introduction

Second-generation ADT, such as abiraterone acetate and enzalutamide, have provided much needed life-extending therapies for recurrent or mCRPC patients. However, the utilization of these more effective ADT therapies has coincided with an increase in the development of AVPC (1). This subset of mCRPC is characterized by poor prognosis and loss of AR signaling (2). The absence of AR signaling in AVPC renders the existing hormone targeting treatments ineffective and remaining approved therapies, including platinum-based chemotherapy, offer only limited therapeutic benefits (3). A subset of AVPC tumors are classified as NEPC because they express neuroendocrine genes which are not typically expressed in prostate adenocarcinoma (AdCa). Recent work has implicated the loss of *RB1* and *TP53* mutations as key alterations in the development of NEPC, and inhibition of kinases such as Aurora A kinase (AURKA), MAPK, or FGFR could provide therapeutic opportunities if selected in the right patient subsets (1,4–6). Even with these new developments, there still remains a critical need to understand the molecular characteristics and kinase signaling pathways of NEPC tumors to identify and validate effective treatment options.

Receptor tyrosine kinases link the extracellular environment to intracellular responses through multiple signaling cascades. These signaling cascades regulate numerous pathways that are frequently altered in transformed cells, including cell growth, metabolism, proliferation, differentiation, invasion, motility, and cell death (7). RET is a receptor tyrosine kinase that is essential for neural crest development and is frequently mutated or translocated in subsets of endocrine tumors such as multiple endocrine neoplasia 2 (MEN2) and papillary thyroid carcinomas, respectively (8). RET can be therapeutically targeted with some success in these tumor types. Recently, RET kinase was identified to be tyrosine phosphorylated in a CRPC patient with small cell neuroendocrine pathology (9) and as an enriched cell surface marker in NEPC (10). Further, RET knockdown restricted invasion and proliferation of prostate adenocarcinoma *in vivo* (11). However, it remains unclear whether RET kinase contributes to the growth of NEPC tumors and whether RET inhibition could be exploited as a therapeutic target in this setting.

Here, we evaluated the phosphoproteome of multiple AR negative and AdCa prostate cancer cell lines to identify altered kinase signaling pathways unique to AR negative prostate cancers. Several downstream signaling networks of RET kinase, and RET kinase itself, were enriched and activated in the AR negative cell lines when compared to AdCa cell lines. Additionally, RET kinase was overexpressed in NEPC tumors in multiple clinical datasets. We evaluated the ability of a novel RET pathway inhibitor, AD80, to block cell growth and reduce cell viability in multiple prostate cancer cell lines and observed that it was more potent than current RET-directed FDA approved therapies, cabozantinib and vandetanib (12,13).

Finally, we found that AD80 was effective in blocking tumor growth and reducing tumor viability of NEPC xenograft tumor models as well as organoid models. These results indicate that RET kinase is active in NEPC, contributes to the survival of NEPC tumors, and inhibiting RET induces cell death in neuroendocrine prostate cancer cells that are resistant to current hormonal therapies. These results ultimately nominate RET as a key candidate to test further in the development and effective treatment of NEPC.

## Results

### AR negative cell lines have altered phosphotyrosine and phosphoserine/threonine kinase signaling pathways

To identify the unique kinase signaling pathways required for growth and proliferation of AR negative, we performed phosphoproteomic profiling to compare androgen responsive, AR-full length positive, AdCa cell lines (LNCaP, VCaP, C4-2, and 22Rv1), to AR low/null cell lines that are resistant to ADT and harbor mutations commonly found in NEPC tumor samples (DU145, PC3, NCI-H660, cMyc/myrAKT, LASCPC-01, EF-1, and PARCB-1,-2,-3, and -5) (Supplemental Table 1). Supervised hierarchical clustering between the AdCa and AR negative groups revealed distinct patterns in phosphoserine/threonine (pS/T) and phosphotyrosine (pY) (Figure 1A and 1B, respectively and Supplemental Tables 2 and 3). Kinase substrate enrichment analysis (KSEA) identified AURKA as the most highly enriched pS/T kinase (Figure 1C) that has been previously reported to be significantly upregulated in NEPC (4). Interestingly, among the tyrosine kinases, RET kinase was also significantly enriched (Figure 1D), suggesting that RET kinase is activated in AVPC cell lines (full pS/T and pY KSEA results are in Supplemental Tables 4 and 5, respectively). We confirmed RET protein to be upregulated in the NEPC subset of AVPC cell lines compared to AdCa and confirmed the loss of full length AR (Supplemental Figure 1). Further investigation into the RET pathway via our cell line-derived and our previously published mCRPC rapid autopsy phosphoproteomic datasets (9) (expanded phosphoproteome data set in Supplemental Table 6, see methods) identified hyper-phosphorylation and, in some cases, activation, of several RET pathway targets including MAPK, AKT, and STAT3 (Figure 1E, 1F), further confirming RET pathway activity in AVPC cell lines and tumors.

**Figure 1.**
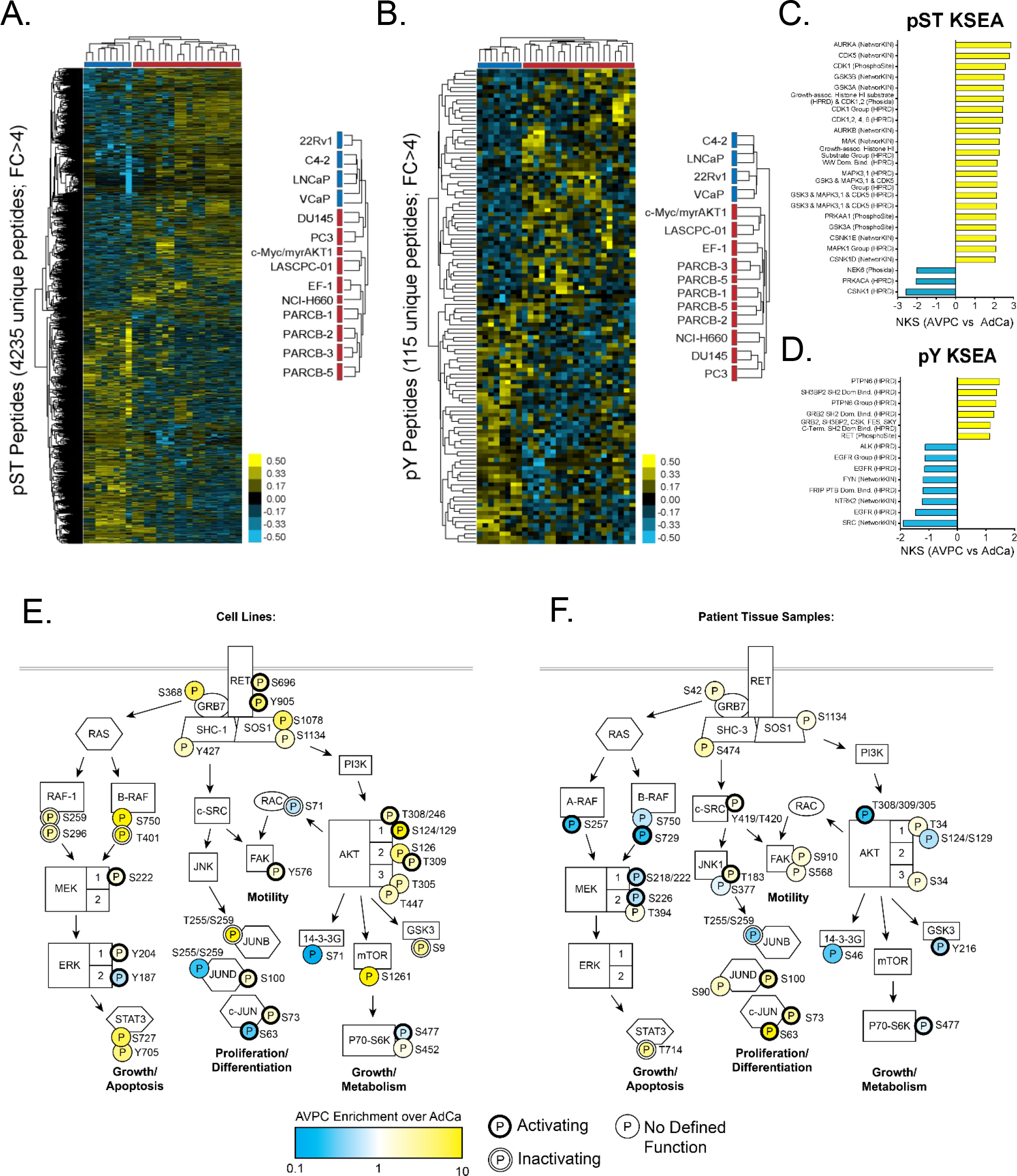
Global phosphorylation and kinase signaling pathways are differentially regulated in AVPC cell lines compared to AdCa cell lines. **A and B.** Supervised hierarchical clustering heatmap of 4,235 unique phosphoserine/threonine (pS/T) enriched peptides (**A**) and 115 unique phosphotyrosine (pY) enriched peptides (**B**) from AdCa cell lines (Blue: C4-2, 22Rv1, LNCaP, and VCaP) and AVPC cell lines (Red: cMyc/myrAKT, LASCPC-01, EF-1, PARCB-1, PARCB-2, PARCB-3, PARCB-5, NCI-H660, DU145, and PC3). Yellow = hyperphosphorylation; Blue = hypophosphorylation. **C and D.** Kinase substrate enrichment analysis (KSEA) performed on the 10 AVPC and 4 AdCa cell lines in A and B, showed multiple alterations to kinase signaling. (**C**) KSEA for pS/T analysis used a false discovery rate (FDR) <0.05, substrate hits > 5, and normalized K score >2.0. (**D**) KSEA for pY analysis used an FDR <0.1, substrate hits >4, and normalized K score >1.1. **E.** Phosphorylated residues identified in the global phosphoproteomics (from **A and B**) or **F.** human phosphoproteome data (19) were mapped onto signaling pathways downstream of RET kinase. Yellow = Enriched in AVPC relative to AdCa; Blue = Reduced in AVPC relative to AdCa. Thick black outline = activating phosphorylation; white outline = inactivating phosphorylation; thin outline = no defined function.

### RET kinase expression is upregulated in patients with neuroendocrine prostate cancer

We took advantage of several clinical prostate cancer gene expression datasets to determine whether RET kinase was overexpressed along with known markers of NEPC. Analysis of the University of Washington rapid autopsy dataset (14) which contains multiple metastatic tumors from CRPC patients revealed that the NEPC (AR negative, neuroendocrine positive) subset had enrichment of RET kinase expression concomitant with increased ASCL1 and chromogranin A (CHGA) and decreased AR, NKX3-1, and KLK3 expression (Figure 2A). Among patients with an AR-negative, NE-positive pathology, there was a strong correlation between levels of RET and ASCL1, while there was no correlation in the other groups (Figure 2B). Additional transcript datasets comparing metastatic NEPC to metastatic AdCa (Figure 2C and Supplemental Figure 2A) (6,15) or LuCaP neuroendocrine prostate cancer patient-derived xenografts (PDXs) with AdCa PDXs (Figure 2D) (16) both showed similar trends of increased RET kinase expression along with upregulation of CHGA and SYP in the NEPC samples. Transcripts from biopsies of progressive, mCRPC samples that followed patients through disease progression revealed clusters of high RET and ASCL1 expression with a strong downregulation of AR regulated genes (Supplemental Figure 2B) (17). Overall, these independent datasets demonstrate that RET kinase is overexpressed in clinical NEPC tumors and support our cell line phosphoproteomic and KSEA analyses, suggesting enhanced RET activity and nominating RET as a candidate therapeutic target for NEPC tumors.

**Figure 2.**
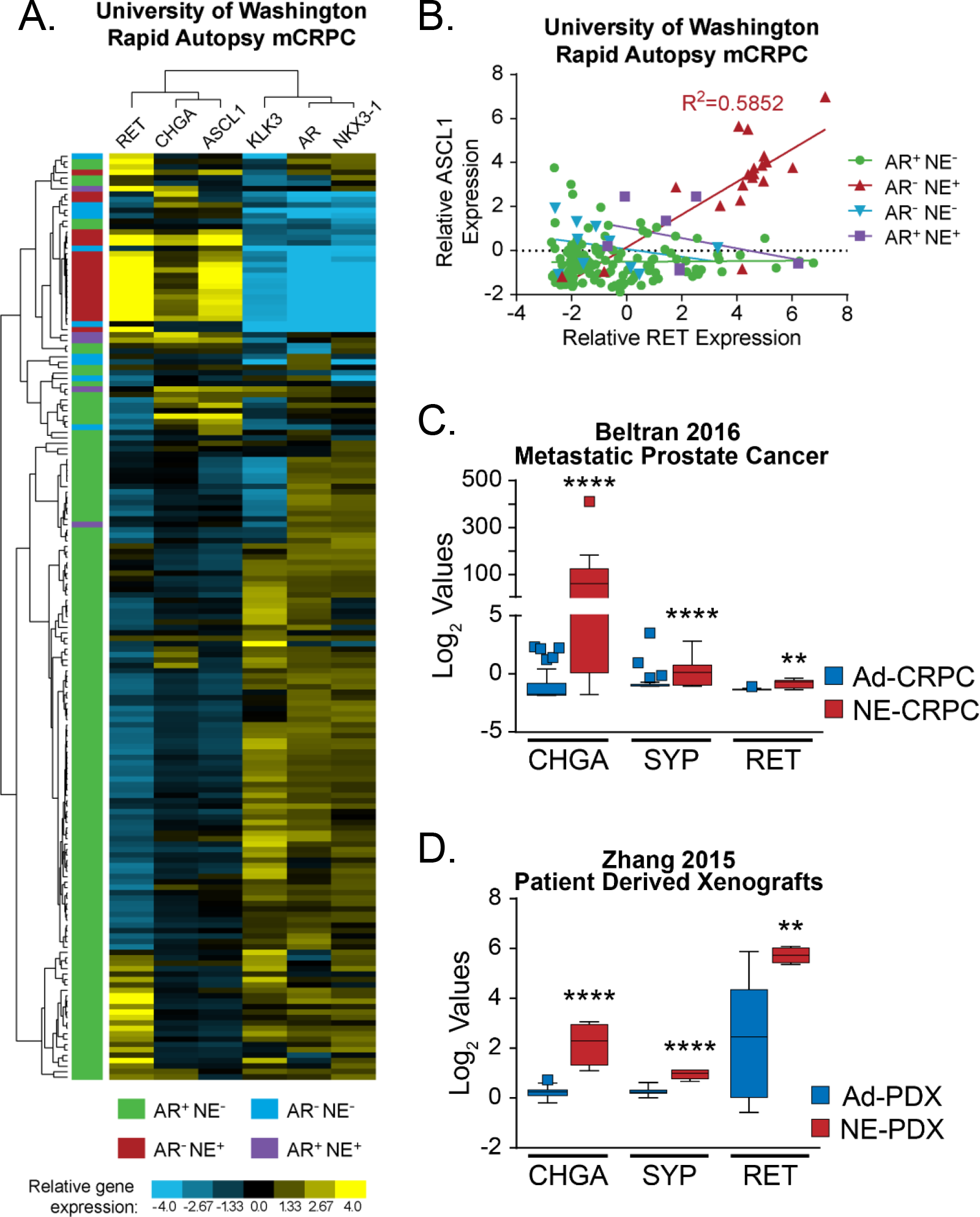
RET kinase along with other neuroendocrine transcripts are upregulated in NEPC relative to AdCa patient samples. **A.** Microarray data from the University of Washington rapid autopsy data of metastatic prostate cancer biopsies published in *Kumar et al. 2016. Nature Medicine.* (14) were clustered based on gene expression of RET, neuroendocrine markers: CHGA and ASCL1, as well as androgen regulated genes: KLK3, AR, and NKX3-1. Upregulation of expression is represented by yellow, while downregulated genes are represented by blue. Patient samples were classified by AR and NE markers as AR+NE-(green, n=134), AR-NE-(blue, n=10), AR-NE+ (red, n=20), and AR+NE+ (purple, n=7). **B.** Scatter plot of relative RET expression values plotted versus relative ASCL1 expression values. Patient samples classifications (as designated in 2A) were plotted individually and analyzed by linear regression. Only the AR-NE+ group had a significant correlation with an R^2^ value of 0.5852. **C.** Whole-exome sequencing of CRPC patient samples from with AdCa (CRPC-Adeno, n=34) or neuroendocrine (CRPC-NE, n=15) phenotype published in *Beltran et al. 2016. Nature Medicine.* (15) show an upregulation of CHGA, SYP, and RET kinase. **D.** Agilent oligo array expression analysis of four neuroendocrine AR-negative LuCaP patient derived xenografts (PDX) and 20 LuCaP adenocarcinoma PDX published in *Zhang et al. 2015. Clinical Cancer Research.* (16) shows an upregulation in CHGA, SYP, and RET kinase. **C and D.** Data represented in Tukey plots and expression values were analyzed by Student’s t test.

### AD80 reduces tumor growth in RET expression high NCI-H660 xenograft tumors by increasing cell death

AD80 is a novel, more selective inhibitor of the RET pathway than previous multi tyrosine kinase inhibitors such as cabozantinib or vandetanib (13). We found that the NEPC cell line, NCI-H660, was among the most sensitive to AD80 treatment (Supplemental Figure 3) and expressed high levels of RET protein (Supplemental Figure 1). To test the effect of AD80 in an *in vivo* model system of NEPC, we generated NCI-H660 xenograft tumors in NOD-SCID mice. Once tumors reached 100-200 mm^3^, mice were randomized and placed into one of three treatment groups: Control (DMSO), 20 mg/kg enzalutamide, or 10 mg/kg AD80 (Figure 3A). Over the course of the 22-day treatment, AD80 treated tumors showed a significant reduction in overall tumor volume (Figure 3B) without a significant effect on animal weight (Figure 3C). This experiment was repeated in a second cohort of mice with 24 days of treatment and higher dose of AD80 (20 mg/kg). The higher dose of AD80 was associated with increased toxicity, but showed similar inhibition of tumor growth throughout the 24-day treatment (Supplemental Figure 4).

**Figure 3.**
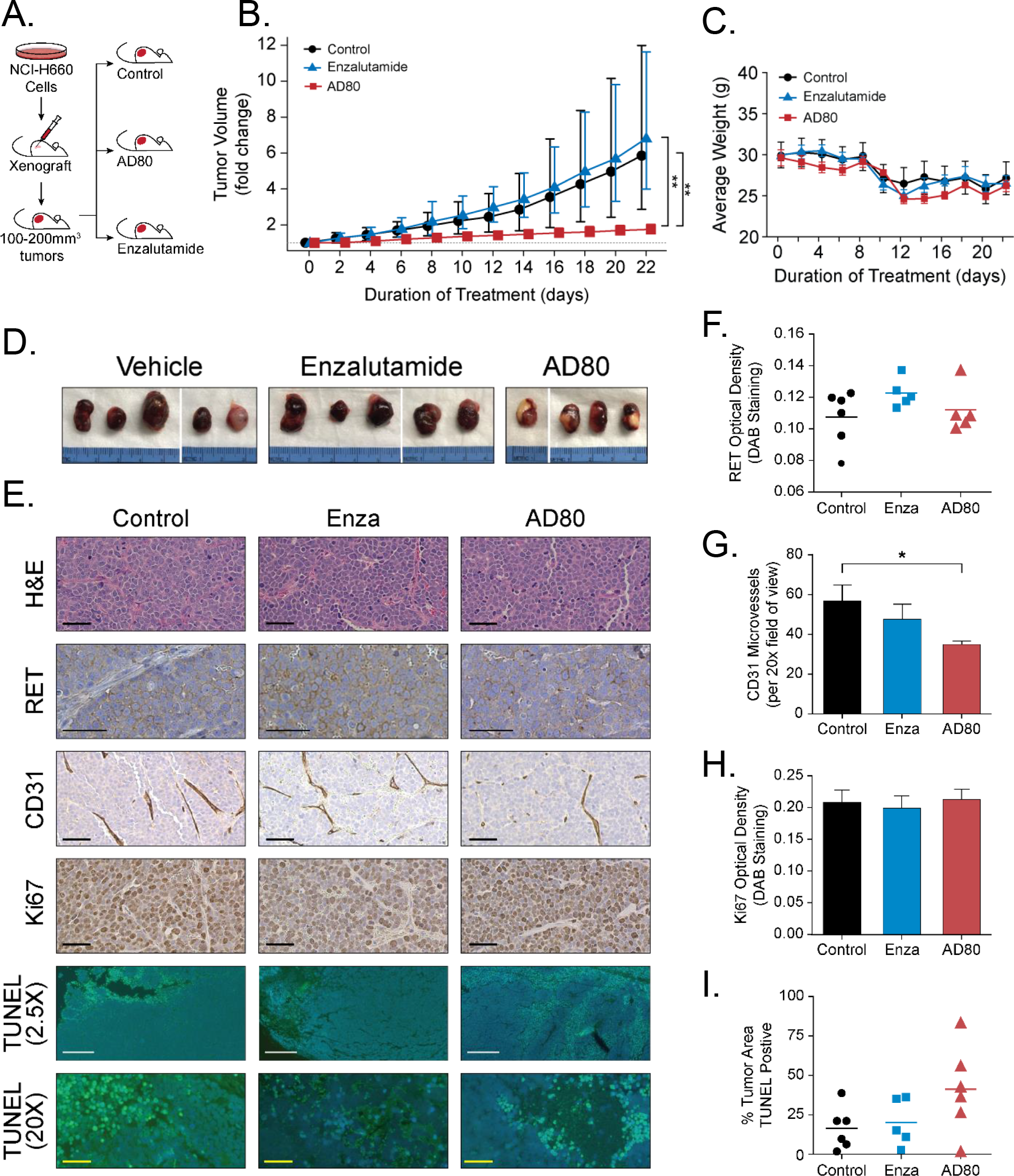
AD80 treatment effectively reduces tumor growth in an NCI-H660 xenograft model by reducing angiogenesis and inducing cell death but has no effect on proliferation. **A.** Schematic of *in vivo* study in which NCI-H660 cells were injected subcutaneously into the right flank of NOD-SCID mice and tumors were allowed to grow to approximately 100 to 200mm^3^ before being randomly assigned into one of three treatment groups: Control (DMSO alone, n=6), ENZA (20mg/kg/day, n=5), or AD80 (10mg/kg/day, n=6). **B.** The fold change in tumor volume by treatment group was plotted as a function of the number of days of treatment. Means and confidence intervals (CIs) were calculated on the log scale and reported in terms of geometric means after exponentiation with error bars ± 95% confidence interval. There was evidence of an overall treatment effect on tumor growth rate (p<0.001) and post-hoc pairwise comparison of the AD80 treated group to DMSO or enzalutamide at day 22 revealed a reduction in mean tumor volume (p<0.01) **C.** Average animal weights were measured at the same time as tumor volumes and no differences in average animal weight between treatment groups was observed over the duration of the study. Symbols represent means with error bars ± standard error. **D.** Following the termination of the study, tumors were excised from animals and photographed with a centimeter scale ruler. Separate images from the same group are divided by a white line. **E.** Representative 20X images of H&E and IHC for total RET, CD31, Ki67, and IF for TUNEL (20x and 2.5X) stained sections of tumors from each experimental arm. Black scale bars are 50μm. White scale bars are 500μm. Yellow scale bars are 50μm. **F-H.** Quantification of the average RET staining intensity (**F**), quantification of CD31 positive microvessels in a field of view (**G**), and average Ki67 staining intensity (**H**) were analyzed by one-way ANOVA. Quantification of the average TUNEL positive area was analyzed with the Kruskal-Wallis test (4X). Symbols represent averages for individual tumors with a horizontal line representing the mean. Bars represent the mean with error bars represent ± standard error.

Following treatment, the AD80 treated tumors appeared smaller and displayed less vascularization (Figure 3D and Supplemental Figure 4C), suggesting that AD80 treatment restricted tumor progression possibly via reduction of angiogenesis which has been reported in other neuroendocrine tumor types treated with RET inhibitors (18). To interrogate the molecular characteristics of the different treatments, the tumors were fixed and sectioned for staining. Sections stained with hematoxylin & eosin (H&E) or IHC staining for RET kinase showed similar tumor morphology and expression and localization of RET kinase (Figure 3E, F and Supplemental Figure 4 D, E). To determine if AD80 treatment reduced angiogenesis, tumor sections from the xenograft tumors were stained with a CD31 antibody (Figure 3E).

The AD80 treated group had a statistically significant reduction in elongated CD31 positive cells compared to the control or enzalutamide treated groups (Figure 3G). The reduction in CD31 staining was not observed in the second cohort of mice, but the tumors were collected at a point when they were growing at similar rate to the control group (Supplemental Figure 4A, D, and F). There was no difference in proliferation as assayed by Ki67 staining among the treatment groups in either cohort of mice (Figure 3E, H, and Supplemental Figure 4D and G). However, TUNEL staining showed large regions of positive staining the percentage of total tumor area that stained positive trended higher in the AD80 treated groups (Figure 3E, I and supplemental Figure 4D and H). Taken together, the staining suggests that AD80 treatment is effective in limiting tumor growth by restricting angiogenesis and inducing death in neuroendocrine cells with high RET expression.

### AD80 induces cell death in NEPC organoid models

We next extended our AD80 treatment to a mouse organoid model of NEPC (5). Tumors derived from the *PTEN*^−/−^*Rb*^−/−^(DKO) mice express higher levels of RET mRNA than *PTEN*^−/−^(SKO) or wild type (WT) animals (Figure 4A) (5). Immunofluorescence staining also confirmed an increase of RET kinase protein in the DKO organoids and low to absent RET kinase in the SKO organoids (Figure 4B). The DKO organoids were resistant to enzalutamide treatment, mimicking the ADT resistant characteristic of NEPC prostate cancer cells that express high levels of RET (Figure 4C). Treating the DKO organoids with increasing concentrations of AD80 induced a dose-dependent increase in cell death, as assayed by live-dead PI staining of the organoids with a calculated LD_50_ of 8.3μM for AD80 (Figure 4D, E). High RET expressing NCI-H660 cells cultured as organoids showed a similar sensitivity to AD80 treatment (Supplemental Figure 5). Thus, inhibiting RET kinase with AD80 in an *in vitro* organoid system of NEPC induced cell death and suggests that the specific population of patients that have high RET expression, are refractory to ADT, and have few remaining therapeutic options, may benefit from RET kinase inhibitor therapies.

**Figure 4.**
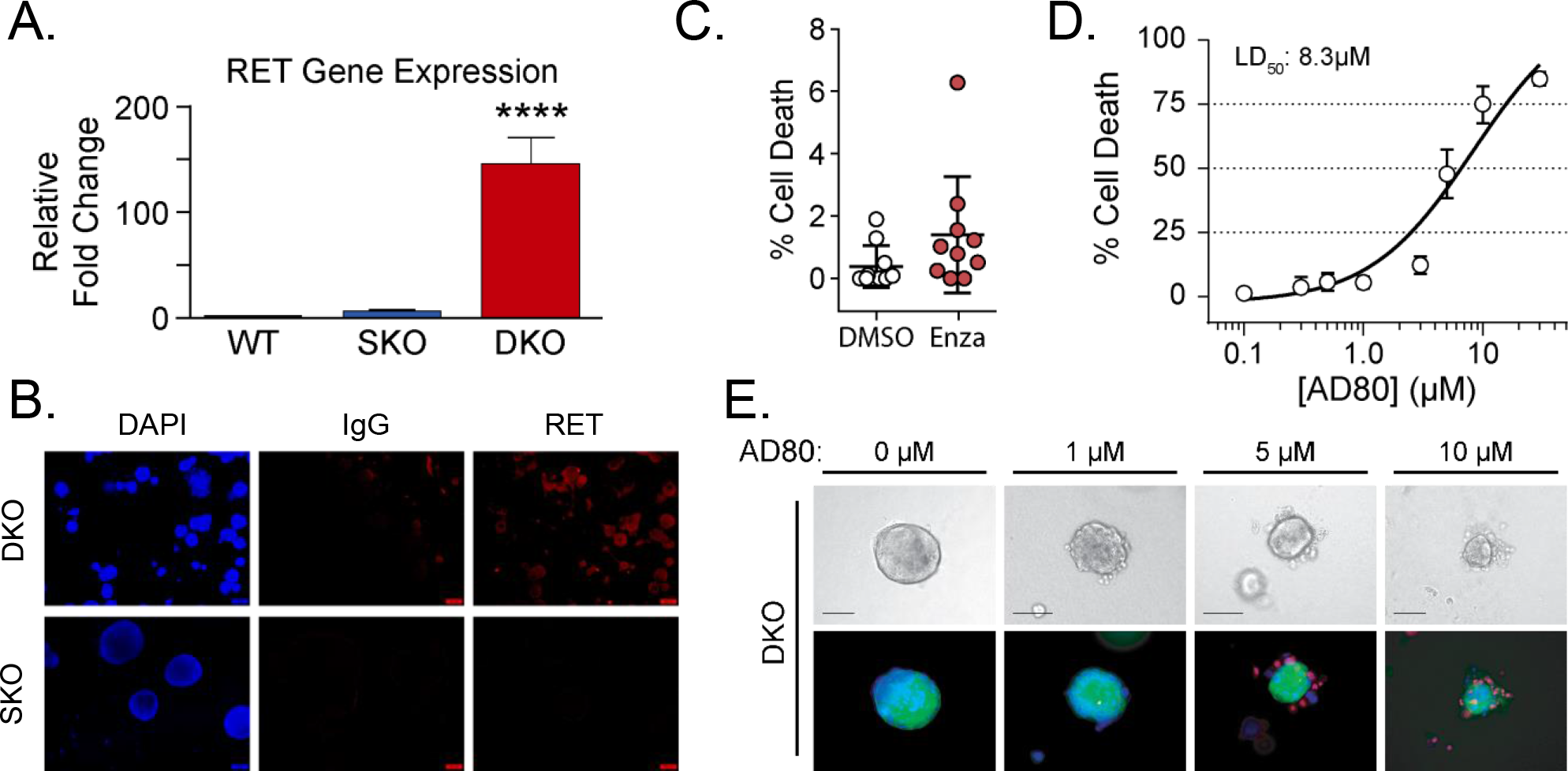
PTEN and Rb KO mouse organoids express high levels of RET and are susceptible to AD80 induced cell death. **A.** RNAseq data of RET gene expression from wild type (WT), PTEN^−/−^ and Rb^−/−^ double knockout (DKO), or PTEN^−/−^ single knockout (SKO) mouse prostate epithelium organoids. RET FKPM values were normalized to WT prostate tissue and expressed as fold change and analyzed by Student’s t test. **B.** RET immunofluorescence staining in PTEN^−/−^ and Rb^−/−^ DKO derived organoids and PTEN^−/−^ SKO organoids or IgG antibody control. DAPI staining of nuclear DNA is used to identify organoids. **C.** Percentage of PI positive cells in DKO organoids treated with DMSO or 10μM enzalutamide. Circles represent values from individual organoids. Horizontal bar represents the mean with error bars ± standard error. **D.** Dose response curve of DKO organoids treated with increasing concentrations of AD80. The LD50 for AD80 alone was calculated to be 8.3μM. Circles represent mean and error bars ± standard error. **E.** Bright field images and corresponding fluorescence images of GFP labeled-DKO organoids treated with the indicated concentrations of AD80. Blue=DAPI staining of nuclei, Red=Propidium iodide staining of dead cells. Scale bar =100μm.

## Discussion

Increasing evidence points to the activation of kinase pathways as possible key mechanisms that bypass AR-targeted therapies and allow the tumors to continue to survive such a harsh therapeutic environment (1,4,19,20). Utilizing phosphoproteomics, we showed that AR-negative cell lines have altered kinase signaling pathways compared to AR-driven adenocarcinomas, which includes activation of RET kinase. Multiple proteins downstream of the RET kinase pathway were phosphorylated on activating residues in both the cell line and in mCRPC autopsy patient samples. RET mutations or activating rearrangements are drivers of tumor development and growth in MEN2, medullary thyroid cancer and small cell and non-small cell lung cancers, and drugs targeting RET can extend survival of these patients (21–23). Cabozantinib, which inhibits RET kinase and other receptor tyrosine kinases including VEGR1/2, has extended survival in certain cancers with activating RET mutations (24,25). In prostate cancer cabozantinib showed promising phase II clinical trials but failed to meet the endpoint criteria in phase III trials (NCT00940225). However, this was tested in a non-stratified patient population and did not focus on NEPC (26). A retrospective evaluation of post-docetaxel patients with CRPC in the COMET-1 and COMET-2 phase III clinical trials where cabozantinib was compared with prednisone and prednisone plus mitoxantrone suggest that a sub population may benefit from cabozantinib treatment, highlighting the importance of molecular stratification of patients for individualized treatments (27–29). Recently, RET knockdown in a prostate AdCa cell line, LNCaP, was reported to restrict tumor growth, but it remains unclear if and how RET contributes to tumor progression in NEPC (11).

We found that overall RET expression in prostate cancer patient samples is highly variable, but that RET kinase expression correlated very strongly with NEPC. Although, there were examples of metastatic and treatment induced NEPC tumors (based on molecular and pathological features) that lack RET gene expression and inversely there were also patients classified as AR-positive adenocarcinomas that had high levels of RET gene expression, but low gene expression of other neuroendocrine markers (Figure 2A and Supplemental Figure 2A). It is important to note that the transition from AdCa to NEPC may be dynamic (5) and RET expression in AR positive tumors may suggest that these tumors are either a heterogeneous phenotype or are transitioning from AdCa to NEPC. Currently, little is known about the regulation of RET gene expression in prostate cancer. Several key epigenetic regulators (such as CBX2, EZH2, BRN2, and SOX2) have been identified as possible modulators that can switch tumors between an AdCa and NEPC state (5,30–32). Alterations in DNA methylation or transcriptional regulation resulting from the loss of proteins such as Rb may further alter RET expression and activity. Therefore, it remains how robust RET expression is gained during the transition from mCRPC to a NEPC phenotype. In small cell lung cancer, ASCL1 was shown to induce RET gene expression and this mechanism of regulation may hold true in NEPC, but has not been validated (33).

Regardless of the dynamics of RET expression in disease progression, we showed that the RET kinase pathway inhibitor AD80 effectively restricted growth in the *Rb/PTEN* knockout organoids and the NCI-H660 cell line and organoids *in vitro*, as well as NCI-H660 tumors *in vivo*. Inhibiting RET kinase induced cell death suggesting that RET kinase signaling may be important for tumor survival. In order to identify patients that could benefit from treatment including RET inhibition, it will be important to generate assays or validate markers of RET activity in NEPC. Pathology, loss of AR signaling, or expression of neuroendocrine genes are not sufficient alone to identify all patients with high levels of RET expression that may benefit from RET targeted therapies. Moving forward, it will be important to identify the subset of patients that would benefit from inhibition of RET kinase. Development of biomarkers for transcriptional activators, RET protein, or markers of RET activity will enable pre-selection of individuals who would benefit from RET inhibitors. Understanding the regulation of RET gene expression, correlation of RET expression and activity and disease progression, as well as the contribution of RET kinase to mCRPC tumor progression could inform better treatment strategies.

## Material and Methods

### Phosphoproteomics of Prostate Cancer Cell Lines

Cultured prostate cancer cells were scraped, pelleted, and snap frozen. Phosphopeptide enrichment and trypsin digestion were performed as previously described (34). Briefly, cells were lysed in 6M guanidium hydrochloride buffer (6M Guanidinium chloride, 100mM Tris pH8.5, 10mM Tris (2-carboxyethyl) phosphine, 40mM 2-chloroacetamide, 2mM Vanadate, 2.5mM Sodium Pyrophosphate, 1mM Beta-glycerophosphate, 10 mg/ml N-octyl-glycoside), sonicated, and cleared. 5mg of total protein was digested with trypsin and a 4G10 antibody-based immunoprecipitation (IP) was used to enrich phosphotyrosine peptides. The IP supernatant containing the phosphoserine/threonine (pS/T) peptides (2.5mg) were de-salted on C18 columns and separated via strong cation exchange chromatography. In separate, parallel reactions the pY and pS/T peptides were enriched from non-phosphorylated peptides using titanium dioxide columns. Finally, the pY and pS/T peptides were each de-salted with C18 tips prior to mass spectrometer analysis (LC-MS/MS with a dual pump nanoRSLC system (Dionex, Sunnyvale CA) interfaced with a Q Exactive HF (ThermoFisher, San Jose, CA) (35)). Technical duplicates were run for all samples and data were analyzed using MaxQuant Andromeda version 1.5.3.30 (parameter settings in (36)) against the Uniprot human reference proteome database with canonical and isoform sequences (downloaded September 2016 from http://uniprot.org). Datasets are accessible through dataset identifiers PXD012970 and PXD012971 (37) through the ProteomeXchange Consortium via the PRIDE partner repository. We expanded upon our previously published mCRPC dataset by decreasing the phosphosite localization probability cutoff from 0.99 to 0.75 (34). This increased our identifications nearly 50% and have now reported those extra identifications in this manuscript as Supplemental Table 6.

Phosphoproteome MS data analysis was performed as previously described (20). For supervised clustering, pY and pS/T data were filtered using a 4-fold change cutoff comparing NEPC vs AdCa from the original excel tables (See Supplemental Tables 2 and 3). Hierarchical clustering was performed using the Cluster version 3.0 with the Pearson correlation and pairwise complete linkage analysis (38). Java TreeView version 1.1.6r4 was used to visualize clustering results (39).

### Kinase Substrate Enrichment Analysis

KSEA was performed as previously described (19). Briefly, phosphopeptides were rank-ordered by average fold change between AR negative (AVPC) vs AR positive (AdCa) prostate cancer cell lines. An enrichment score was calculated using the Kolmogorov-Smirnov statistic and statistical significance was calculated via permutation analysis. The normalized enrichment score (NES) was calculated by taking the enrichment score and dividing by the mean of the absolute values of all enrichment scores from the permutation analysis. The Benjamini-Hochberg procedure was utilized to calculate false discovery rate for each kinase. For pY analyses, cutoffs of FDR<0.05, hits>4, and NES>1.3 were used. For pS/T analyses, cutoffs of FDR<0.02, hits>5, and NES>2 were used.

### Tissue Culture

Human prostate cancer cell lines LNCaP, VCaP, C4-2, 22Rv1, DU-145, PC3 and NCI-H660 cells were obtained from ATCC. LNCaP, VCaP, C4-2, 22Rv1, DU145, and PC3 cells were grown in appropriate media as recommended by ATCC (Life Technologies) supplemented with 10% fetal bovine serum (Sigma Aldrich) and 1% penicillin-streptomycin (Life Technologies). NCI-H660 cells were grown in RPMI-HITES medium: RPMI, 5% FBS, 10 nM hydrocortisone (Sigma), 10 nM beta-estradiol (Sigma), insulin-transferrin-selenium (Life Technologies), 1% penicillin-streptomycin, and Glutamax (Life Technologies). LASCPC-01, cMyc/myrAKT, PARCB2 and PARCB5, and EF1 cell lines were obtained from Dr. Owen Witte at UCLA and cultured as described (10,40,41). H660 organoids were cultured as described in (42). LNCaP95 cells were cultured in RPMI 1640 with no phenol red supplemented with 10% charcoal-stripped serum, Glutamax, and 1% penicillin-streptomycin. Mouse organoids were established by enzymatic digestion of GEMM primary prostate tumor tissue in 5 mg/ml Collagenase type II (Gibco) in adDMEM/F12 (Gibco) media with 10 μM Y-27632 dihydrochloride (Tocris Bioscience). Digested cells were seeded into 100% Matrigel and cultured as described by Drost et. al 2016. NCI-H660 organoids were seeded into Prostate 18 QGel 3D Matrix (QGel) according to manufacturer’s instructions and cultured in RPMI-HITES media with B27 supplement (Gibco), 1.25 mM N-acetylcysteine (Sigma), 5 ng/mL EGF (PeproTech), 500 nM A83-01 (Tocris Bioscience), 5 ng/mL FGF2 (PeproTech), 10 ng/mL FGF10 (PeproTech), 10 mM Nicotinamide (Sigma), and 1 μM Prostaglandin E2 (Tocris Bioscience). Culture media was replenished every 4 days and organoids were passaged by sequential digestion in 1 mg/mL Dispase II (Gibco) followed by TrypLE Express (Gibco) and mechanical disruption through a needle to dissociate to single cells before re-suspension as a 3D culture. All cell lines were grown and maintained in a humidified incubator at 37°C and 5% CO_2_.

### Antibodies

The following antibodies were used for IHC and IF: RET (Cell Signaling Technology, E1N8X, IF: 1:100 IHC 1:500), CD31 (Cell Signaling Technology, D8V9E, 1:100), and Ki67 (Cell Signaling Technology, D2H10 1:400).

### In vivo study

Experiments were carried out on 8 week old male NOD-SCID mice in accordance with IACUC approved protocols. Xenografts were generated via subcutaneous injection of 1×10^6^ NCI-H660 cells per animal mixed at a 1:1 ratio with Corning Matrigel Matrix into the right flank. Tumors were allowed to grow to approximately 100-200mm^3^ before mice were randomly allocated into vehicle (5% DMSO), ENZA (20mg/kg/day), and AD80 (10 mg/kg/day or 20mg/kg/day) treatment groups. Treatment proceeded once daily, 5 days a week for 22 days by oral gavage. Tumor volume and animal weight were measured every two days. Tumors volume was measured by caliper and expressed in mm^3^ (Tumor Volume = 0.5 a × b^2^, where a and b represents long and short diameter respectively).

### Immunohistochemistry

Xenograft tumors were formalin fixed paraffin embedded and sectioned following standard procedure. To stain, sections were deparaffinized by baking at 65°C for one hour and hydrated with sequential washes in xylenes, 100% ethanol, 95% ethanol, 70% ethanol and 1×PBS, prior to citrate buffer pH 6.0 antigen retrieval. To stain, tissues were washed with 0.1%TBST, blocked with 2.5% normal horse serum for one hour at room temperature before incubating in primary antibody overnight at 4°C in a humidified slide box. Slides were washed with 0.1%TBST and incubated in HRP-conjugated secondary antibody (Vector Laboratories, MP-7500-15) for one hour at room temperature and developed using a DAB peroxidase substrate kit (Vector Laboratories, NC9567138). Reaction was stopped with water before proceeding to counterstaining with hematoxylin for one minute. Slides were de-stained in tap water, dehydrated with ethanol and xylenes and mounted.

### Terminal deoxynucleotidyl transferase dUTP nick end labeling (TUNEL) assay

The Click-iT™ Plus TUNEL Assay for In Situ Apoptosis Detection, Alexa Fluor™ 488 Kit was used according to the manufacturer’s protocol (Invitrogen). Nuclei were counterstained with Hoechst 33342 (ThermoFisher). A DNase treated positive control section was incubated in 1 U of DNase I diluted into 1X DNase I Reaction Buffer (20 mM Tris-HCl, pH 8.4, 2 mM MgCl_2_, 50 mM KCl) for 30 minutes at room temperature (Invitrogen). The TUNEL‑positive cells in tissue sample slides were identified by comparing to the DNase treated positive control and the no-TdT enzyme negative control.

### Image analysis and quantification of immunohistochemistry

Tumor sections were imaged on a Zeiss Axiovert A2. Average RET or KI67 staining was determined by color deconvolution followed by measurement of the mean gray value in the DAB channel in Fiji (43). Mean gray value was converted to optical density with the following equation: OD=Log(Max gray value/Mean gray value). Values for images from five distinct fields of view were averaged to create a single data point for each tumor in each tumor group. CD31 positive microvessels were identified after color deconvolution and analysis of particles greater than 200 pixels with a circularity between 0.00-0.45 in Fiji. Values from 4-10 distinct 10x fields of healthy CD31 stained tissue were analyzed and averaged to create a value for each tumor in each treatment group. Percent TUNEL positive area was determined by using Fiji to measure the TUNEL positive area divided by total tumor area x100 for each tumor.

### Organoid dose response

For assays, organoids were seeded as single cells in 40 μL of 33% Matrigel (mouse organoids) or Prostate 18 QGel 3D Matrix (NCI-H660 organoids) in 96-well tissue culture plates and cultured for 2 days at 37°C to allow organoid formation. Once formed, organoids were treated with AD80 (at concentrations of ranging from 0.1 μM to 30 μM), or DMSO, with 10 μM Enzalutamide (MedchemExpress) for 72 hours. After treatment, cells were stained with 10 μL ReadyProbes Cell Viability Imaging Kit Blue/Red (Invitrogen) per well for 30 minutes at room temperature and z-stack images of stained cells were taken using an EVOS FL Auto 2 Cell Imaging System (Invitrogen). The percentage of cell death was calculated by identifying the percentage of PI-positive cells per organoid in at least 10 organoids for each treatment condition.

### Statistical Analysis

For xenograft tumor volume experiments, means and confidence intervals (CIs) were calculated on the log scale due to skew and reported in terms of geometric means after exponentiation. Geometric mean tumor volumes were compared on the final day by treatment group using a linear regression model adjusted for baseline log tumor volume. When there was a significant overall treatment effect, post-hoc pairwise comparisons of the treatment groups were made with p-values adjusted using the multcomp package in R. All other statistical analyses were performed using GraphPad Prism 7 with the tests indicated in the figure legends. P<0.05 was considered to indicate a statistically significant difference. P values were determined with significance indicated as follows; *p<0.05, **p<0.01, ***p<0.001 and ****p<0.0001.

## Acknowledgements

We thank members of the Drake lab for providing advice and input on the manuscript. We also thank the members of the Biological Mass Spectrometry Facility of Robert Wood Johnson Medical School and Rutgers, The State University of New Jersey, for providing advice and performing mass spectrometry on our samples. We thank Ryder Clifford from QGel for providing reagents. We thank the patients and their families, Celestia Higano, Evan Yu, Elahe Mostaghel, Heather Cheng, Bruce Montgomery, Mike Schweizer, Andrew Hsieh, Daniel Lin, Funda Vakar-Lopez, Lawrence True and the rapid autopsy teams for their contributions to the University of Washington Medical Center Prostate Cancer Donor Rapid Autopsy Program and the Development of the LuCaP PDX models. This work was supported by the Department of Defense grant (W81XWH-12-1-0399), the Prostate Cancer Biorepository Network (PCBN) (W81XWH-14-2-0183), the Pacific Northwest Prostate Cancer SPORE (P50CA97186), the PO1 NIH grant (PO1 CA163227), the Richard M. LUCAS Foundation, and the Institute for Prostate Cancer Research (IPCR). LCC and VT are supported by the National Institute of General Medical Sciences of the National Institutes of Health under award number T32 GM008339. JMD is supported by the Department of Defense Prostate Cancer Research Program W81XWH-15-1-0236 and W81XWH-18-1-0542, Prostate Cancer Foundation Young Investigator Award, and the New Jersey Health Foundation. Research reported in this publication was supported by the National Center for Advancing Translational Sciences of the National Institutes of Health Award Number UL1-TR002494, NIH Award P50 CA097186, DOD Award W81XWH-18-0347. The content is solely the responsibility of the authors and does not necessarily represent the official views of the National Institutes of Health.

## Supplemental Materials and Methods

### Western Blot Analysis

The following antibodies were used for western blot analysis: Total RET (Cell Signaling Technologies E1N8X, 1:1000), AR (Santa Cruz sc-7305, 1:500) and α-Tubulin (Santa Cruz sc32233, 1:1000). Cells were lysed with 1% SDS/2% β-ME and boiled for 10 minutes following a freeze thaw after lysis. The protein concentration was determined using BioRad Quick Start Bradford Protein Assay Kit following manufacturer’s protocol. 20ug of protein per lane was loaded into GenScript SurePage 4-12% gel, transferred to a nitrocellulose membrane, blocked in 5% BSA in 1×TBST for one hour at room temperature before incubating in primary antibodies (diluted in block) overnight at 4°C. Membranes were washed with 1×TBST before incubating in Licor IR-conjugated secondary antibodies (diluted 1:5000 in block) for one hour at room temperature, washed again and imaged using the Licor Odyssey System and adjusted with the Licor Image Studio Lite software (v5.2).

### IC_50_ value measurement

Cell viability was measured using the WST reagent (Takara) following manufacturer’s protocol. Cabozantinib, vandetanib and AD80 were all obtained from Selleckchem. All compounds were tested against human prostate cancer cell lines: LNCaP, DU145, C4-2, 22Rv1, PC3, NCI-H660. Only AD80 was tested against VCaP, LNCaP95, and EF1 cell. Cell densities were determined for 96-well plates prior to performing the assay. Cells were treated with drug for 72 hours prior to the WST assay. Each concentration data point was conducted in triplicate. Each compound was tested at a minimum of ten dose levels, separated by three-fold dilution concentration intervals, IC_50_ values were calculated using GraphPad Prism 7. Reported values were calculated from a single WST assay, but were confirmed by repeating the entire assay in duplicate.

## Supplemental Figure Legends

**Supplemental Figure 1.**
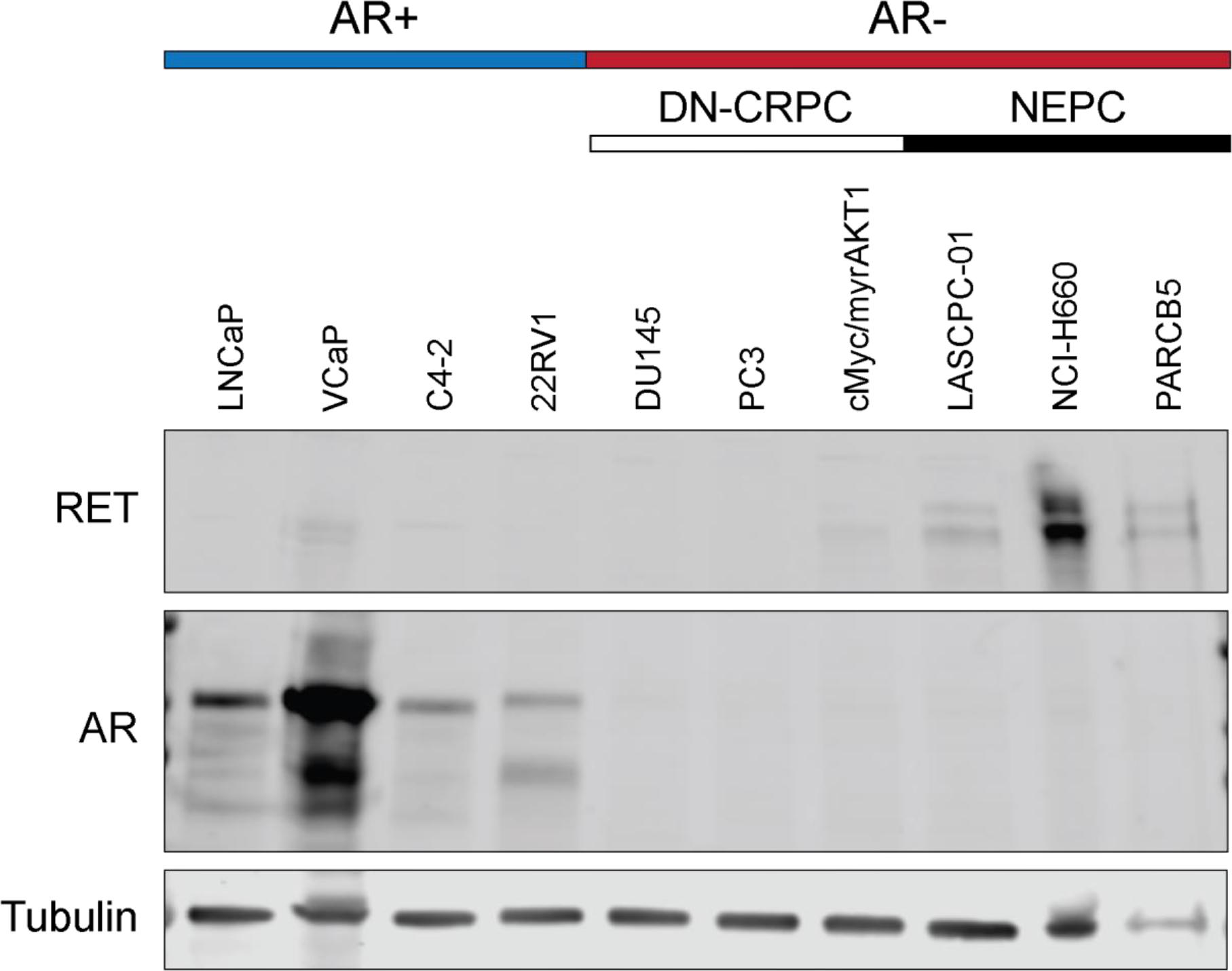
RET expression is higher in prostate cancer lines that have lost androgen receptor signaling. Cell lines used in the global phosphoproteomic analysis were analyzed by western blot for expression of total RET kinase, AR, and α-Tubulin as a loading control. LNCaP, VCaP, C4-2, and 22Rv1 cells were classified as AR-positive (blue bar), while the remaining cell lines were AR-negative (red bar). The AR-negative cell lines can further be broken down into double negative castration resistant prostate cancer (DN-CRPC, white bar) or neuroendocrine prostate cancer (NEPC, black bar).

**Supplemental Figure 2.**
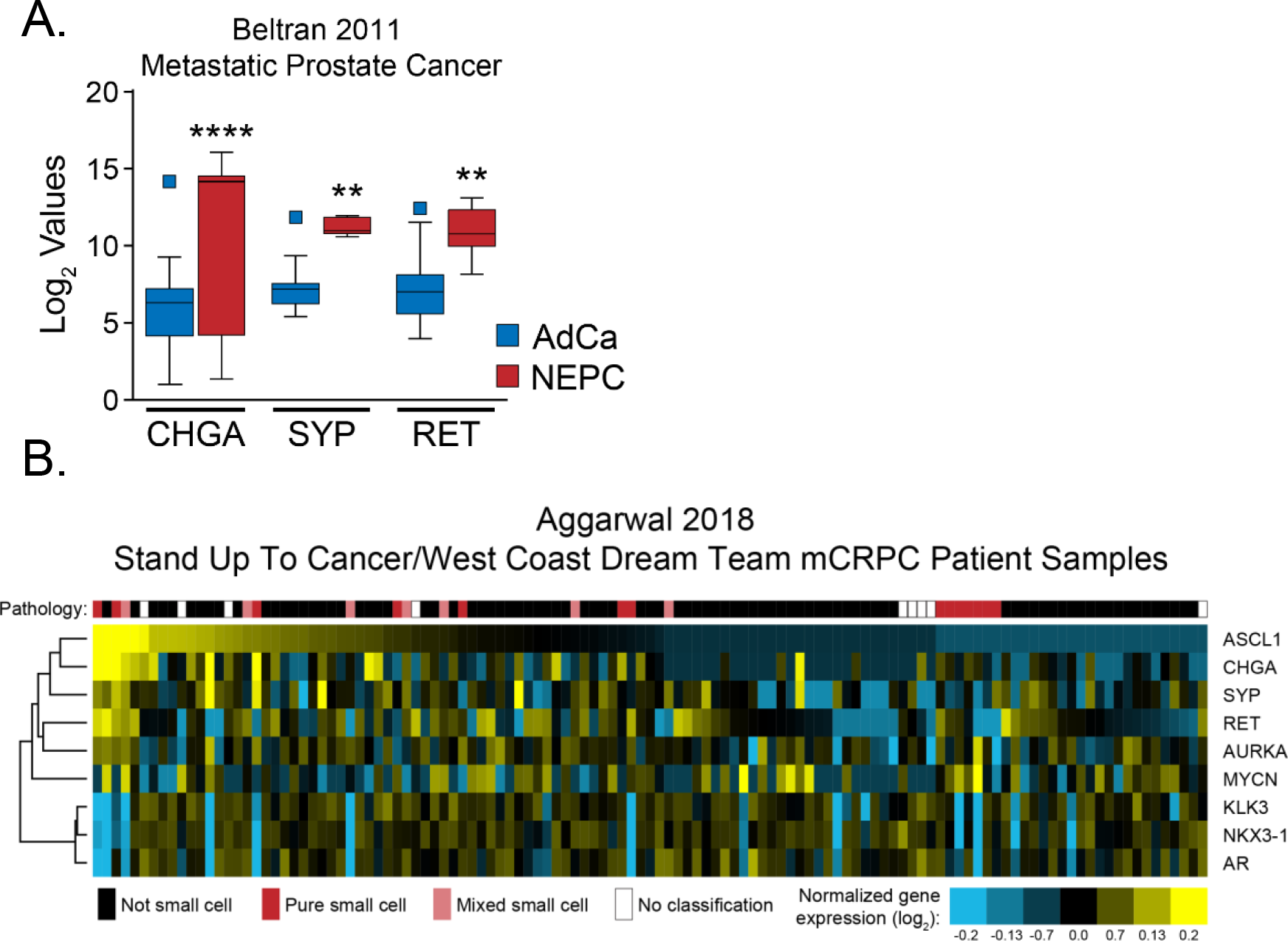
RET gene expression is higher in patient samples with pure/mixed phenotype or primary NEPC samples, along with other neuroendocrine markers. **A.** Next generation sequencing data of 30 adenocarcinoma and seven NEPC patient samples from *Beltran et al. 2011. Cancer Discov.* (6) shows an upregulation in neuroendocrine markers chromogranin A (CHGA) and synaptophysin (SYP) and also RET kinase. **B.** mCRPC patient samples collected by the Stand Up To Cancer/West Coast Dream team (17) were sorted by ASCL1 gene expression and clustered based on selected neuroendocrine and AR responsive genes. Yellow represents upregulated genes, while blue represents downregulated genes. Pathology classifications are designated by colored squares above patient samples (Black: not small cell; Red: Pure small cell; Pink: Mixed small cell; White: no pathology classification).

**Supplemental Figure 3.**
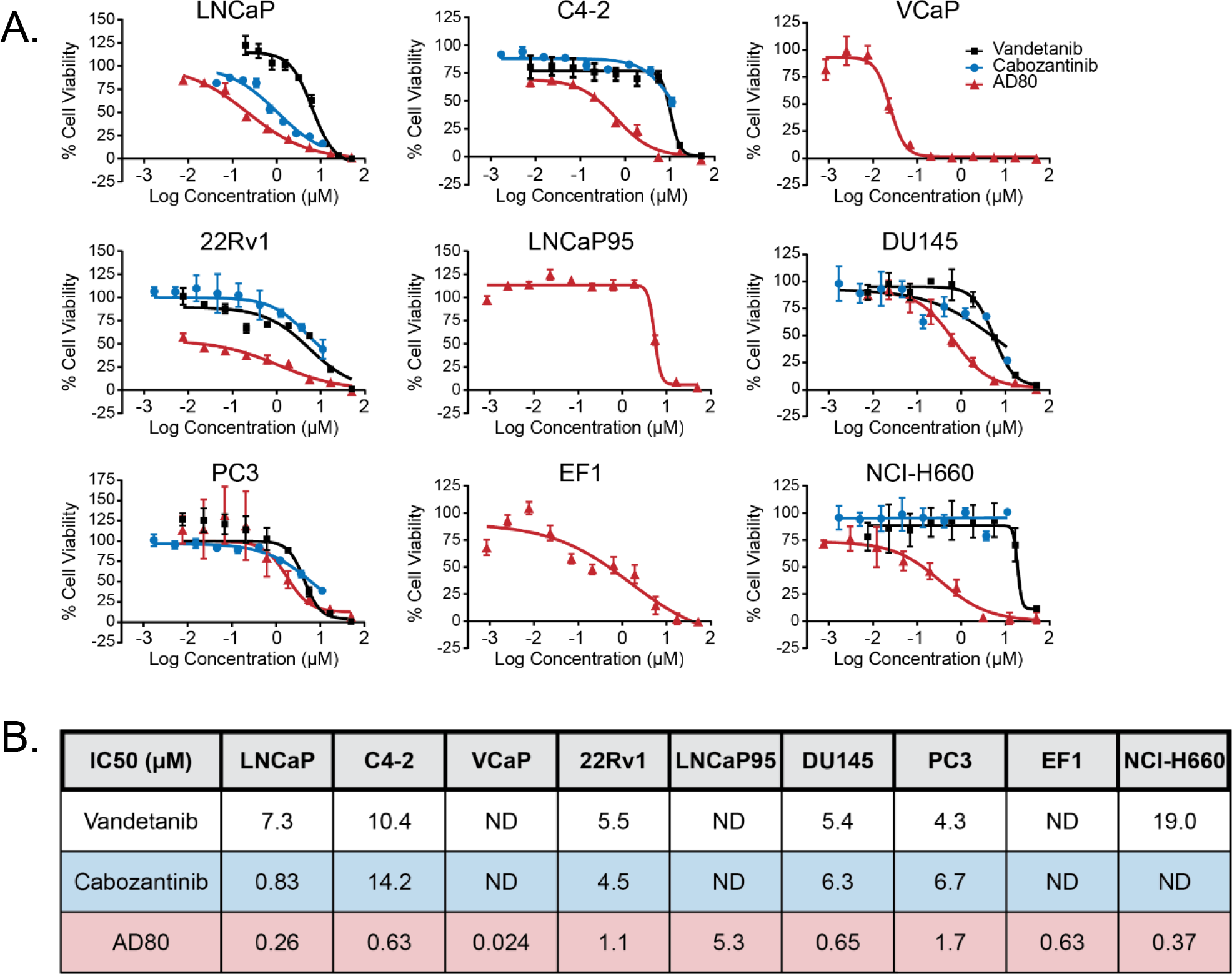
NEPC cells show greater relative sensitivity to AD80, a potent RET inhibitor, compared to other multi kinase inhibitors than AdCa or AVPC cell lines. **A.** IC_50_ dose response curves of AdCa (LNCaP, C4-2, and 22Rv1), AVPC (DU145 and PC3), and NEPC (NCI-H660) cell lines treated with varying concentrations of multi-tyrosine kinase inhibitors vandetanib, cabozantinib, AD80. Error bars represent ± SD. **B.** Table of the calculated IC_50_ values (μM) for each of the cell lines and three drugs. ND is not determined.

**Supplemental Figure 4.**
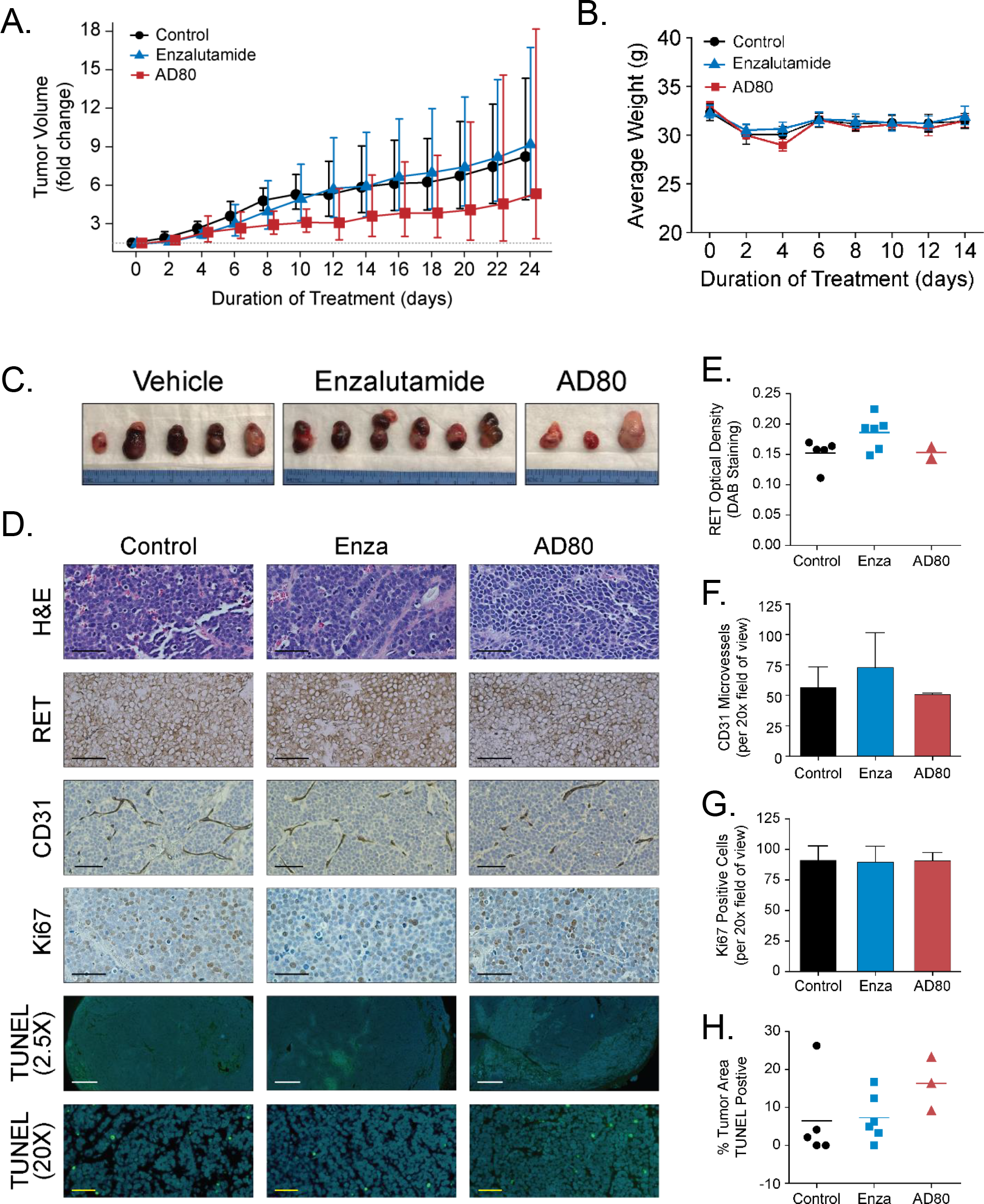
A higher dose replicate of AD80 treatment of NCI-H660 xenograft tumors shows a similar effect of restricting tumor growth. Animals were assigned to the following treatment groups: Control (DMSO alone, n=5), ENZA (20mg/kg/day, n=6), and AD80 (20mg/kg/day, n=3). **A.** The fold change in tumor volume by treatment group was plotted as a function of the number of days of treatment. Means and confidence intervals (CIs) were calculated on the log scale and reported in terms of geometric means after exponentiation. Error bars ± 95% confidence interval. Over the 24 day treatment course AD80 reduced the overall growth rate of the tumors (p<0.05) but there was no difference in the mean tumor volumes at the 24 day time point. **B.** Animal weights were measured at the same time as tumor volumes and represented as an average for the treatment group. Symbols represent mean with error bars ± standard error. **C.** Following the termination of the study, tumors were excised and photographed with a 1cm scale ruler. **D.** Representative 20X images of H&E and IHC for total RET, CD31, Ki67, and IF for TUNEL (20x and 2.5X) stained sections of tumors from each experimental arm. Black scale bars are 50μm. White scale bars are 500μm. Yellow scale bars are 50μm. **E-H.** Quantification of the average RET staining intensity (**E**), quantification of CD31 positive microvessels in a field of view (**G**), and average number of Ki67 positive stained cells from 3 distinct 20X fields of view for each tumor (**H**) were analyzed by one-way ANOVA. Quantification of the average TUNEL positive area was analyzed with the Kruskal-Wallis test (4X). Symbols represent averages for individual tumors with a horizontal line representing the mean. Bars represent the mean with error bars ± standard error.

**Supplemental Figure 5.**
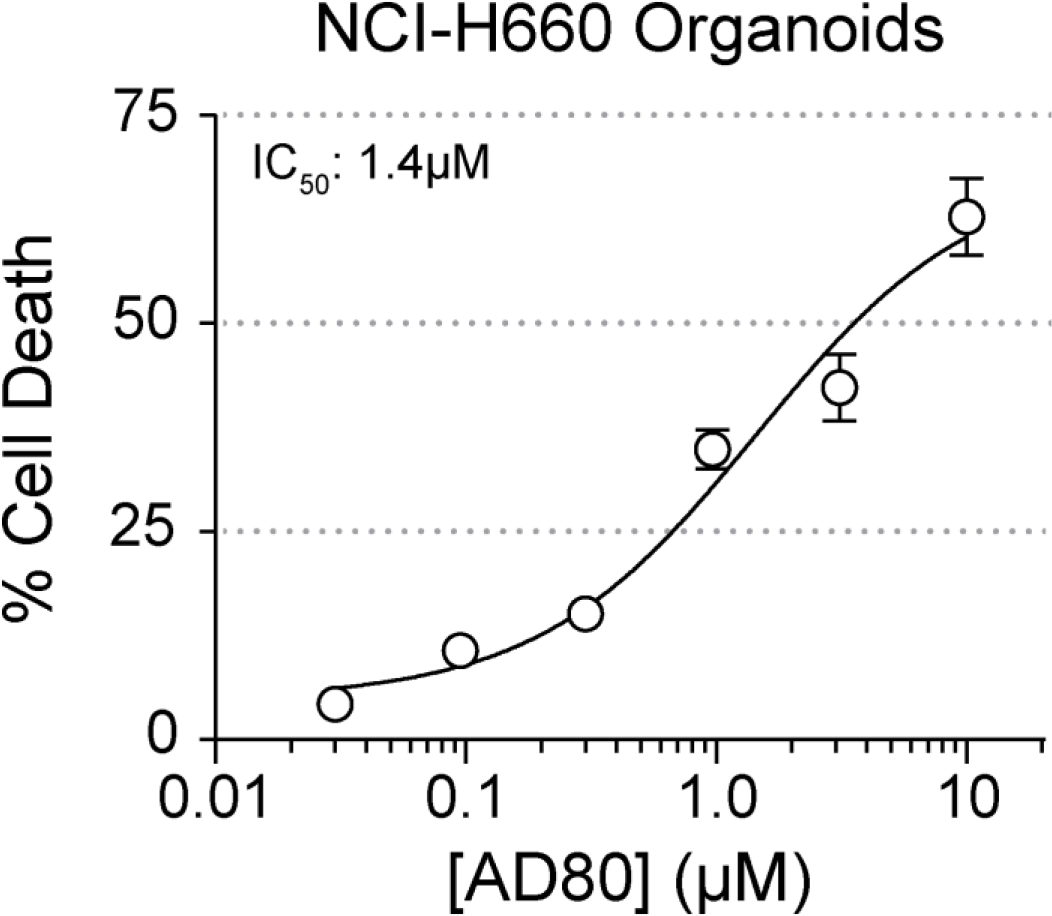
AD80 treatment induces cell death in NCI-H660 cells cultured as organoids. Dose response curve of DKO organoids treated with increasing concentrations of AD80. The IC50 for AD80 was calculated to be 1.4μM. Circles represent the mean with error bars ± standard error.

